# Defective transfer of parental histone decreases frequency of homologous recombination in budding yeast

**DOI:** 10.1101/2023.01.10.523501

**Authors:** Srinivasu Karri, Yi Yang, Jiaqi Zhou, Quinn Dickson, Zhiquan Wang, Haiyun Gan, Chuanhe Yu

## Abstract

Recycling of parental histones is an important step in epigenetic inheritance. During DNA replication, DNA polymerase epsilon subunit DPB3/DPB4 and DNA replication helicase subunit MCM2 are involved in the transfer of parental histones to the leading and lagging DNA strands, respectively. Single *Dpb3* deletion (*dpb3Δ*) or *Mcm2* mutation (*mcm2-3A*), which each disrupt one parental histone transfer pathway, leads to the other’s predominance. However, the impact of the two histone transfer pathways on chromatin structure and DNA repair remains elusive. In this study, we used budding yeast *Saccharomyces cerevisiae* to determine the genetic and epigenetic outcomes from disruption of parental histone H3-H4 tetramer transfer. We found that a *dpb3Δ*/*mcm2-3A* double mutant did not exhibit the single *dpb3Δ* and *mcm2-3A* mutants’ asymmetric parental histone patterns, suggesting that the processes by which parental histones are transferred to the leading and lagging strands are independent. Surprisingly, the frequency of homologous recombination was significantly lower in *dpb3Δ, mcm2-3A*, and *dpb3Δ*/*mcm2-3A* mutants relative to the wild-type strain, likely due to the elevated levels of free histones detected in the mutant cells. Together, these findings indicate that proper transfer of parental histones to the leading and lagging strands during DNA replication is essential for maintaining chromatin structure and that high levels of free histones due to parental histone transfer defects are detrimental to cells.

## Introduction

In eukaryotes, genomic DNA is packaged into an organized structure called chromatin. Each nucleosome core particle, which represents the basic unit of chromatin, is composed of two copies of each of the histones H2A, H2B, H3, and H4. Theses histones are assembled into an octameric core and tightly wrapped around a section of 147 bp of DNA (1). Upon the arrival of the DNA replication fork, the core histone octamer is disrupted; following the DNA replication fork, the histone octamer is reassembled with both parental histones and newly synthesized histones (2,3). These recycling of each type of parental histone molecules behaves differently during DNA replication. The histone H3-H4 tetramers do not segregate and are transferred to new DNA (4). The recycle of histones H2A/H2B occurs independently of H3-H4 tetramer; and H2A-H2B dimers dissociate from the parental octamer and randomly reincorporate into newly assembled nucleosomes (5,6). The transfer of parental histone H3-H4 tetramers during chromatin replication is the essential first step in epigenetic inheritance (2,7). During every cell cycle, cells synthesize half the required histone molecules for DNA replication, with the other half coming from recycled parental histones (2,3). In yeast, the amount of parental H3-H4 tetramers transferred to the leading and lagging strands is nearly equal, though a slight bias exists for the lagging strand (8,9). DNA Polymerase ε subunit *Dpb3/Dpb4* mediates parental histone H3-H4 tetramer transfer to the leading strand (9). The *Mcm2-Ctf4-Polα* axis facilitates transfer of parental H3-H4 tetramers to the lagging strand. Specifically, MCM2, a subunit of the replicative helicase CMG, contains a histone-binding motif (HBM) that plays an essential role in mediating parental histone H3-H4 tetramer transfer (8,10,11). Mutations in the HBM, *Ctf4*, or *Polα*/primase that disrupt the CMG helicase’s ability to connect to POLα, lead to a defect in transfer of the parental histone H3-H4 tetramer to the lagging strand (8,11). Defects in the transfer of parental H3-H4 tetramers to either the leading or lagging strands compromise silencing at the *HML* locus in yeast and dysregulate gene expression in mouse embryonic stem cells (12,13). However, the impact of disrupting specific pathways for parental histone transfer on chromatin structure and DNA repair remains elusive.

Homologous recombination (HR) is an error-free mechanism for the DNA damage repair pathway, which is essential to maintain genome stability. Previous studies have shown that HR is regulated by the chromatin environment (14). Several new histone H3-H4 chaperones are involved in this regulation. The chaperone CAF1 is conserved among all eukaryotes. It binds to the H3-H4 tetramer and mediates replication-coupled nucleosome assembly through interaction with PCNA (15,16). *Caf1* mutants show increased homologous recombination in plant likely due to increased DNA accessibility (17). But in human cell lines, immunofluorescence staining showed that CAF1 is locally recruited to DNA double strand break sits (18). The chaperone ASF1 presents the newly synthesized histone H3-H4 to Rtt109-Vps75 for acetylation of histone H3K56, after which the ASF1-bound H3-H4 is transferred to CAF1 or RTT106 (19,20). CAF1 and ASF1 promote homologous recombination through nucleosome assembly (21,22). These studies suggest a connection between the new histone chaperones and HR. More recently, the parental histone deposition pathways have been implicated in template switch recombination, an error-prone of HR, in yeast(23). It is still unclear whether there is a connection between parental histone chaperones and HR.

In this study, we compared a *dpb3Δ*/*mcm2-3A* double mutant to single *Dpb3* and *Mcm2* mutants, as well as the wild-type (WT) strain, in experiments characterizing the strand bias of parental histone transfer, chromatin structure, genomic instability, and HR. We show that the leading and lagging strand parental histone transfer process are independent as the double mutant (*dpb3Δ*/*mcm2-3A*) neutralizes the single mutant’s asymmetric parental histone pattern. The homologous recombination frequency is significantly decreased in *mcm2-3A, dpb3Δ*, and the double mutants, likely due to elevated free histone level. Our findings provide evidence that proper transfer of parental histones to the leading and lagging strands during DNA replication is required for maintaining chromatin structure and genome integrity.

## Results

### A symmetric distribution of parental histones H3-H4 at replicating DNA strands is detected in *dpb3Δ*/*mcm2-3A* double mutant cells

To detect whether interference can take place between the two nucleosome assembly pathways, we first generated a double mutant strain *dpb3Δ*/*mcm2-3A*. We then used enrichment and sequencing of protein-associated nascent DNA (eSPAN) to analyze the binding of parental and newly synthesized H3-H4 tetramers to leading and lagging DNA strands in the *dpb3Δ*/*mcm2-3A* strain. Hydroxyurea was used to arrest synchronized cells in early S-phase (**Fig 1A**) (24), after which H3K4me3 was used to track parental histone H3, and H3K56Ac was used to track newly synthesized histone H3. eSPAN mapping snapshots are displayed at **Fig 1B** and **Suppl Fig1B**.

**Figure 1.**
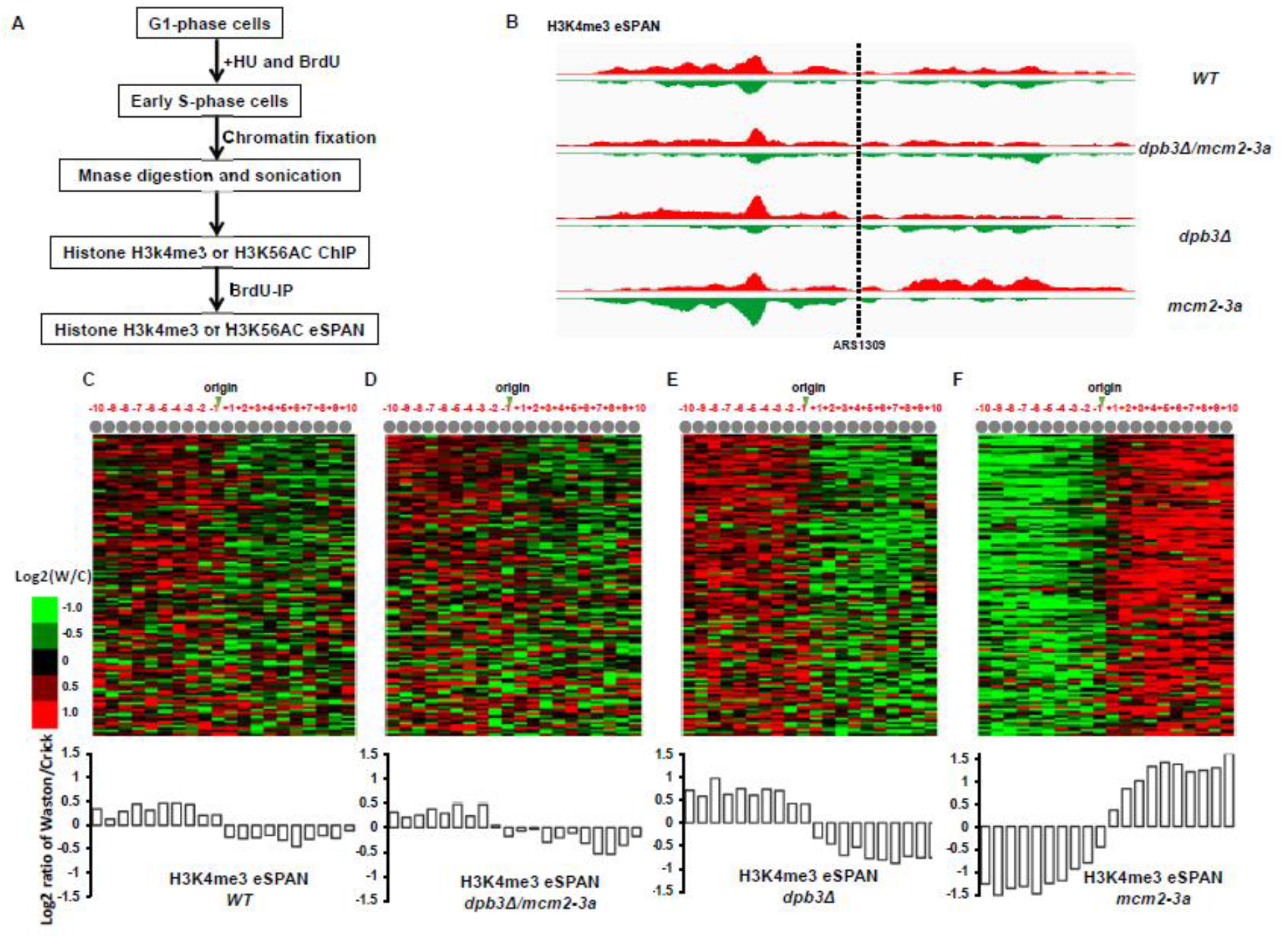
Combination of *dpb3Δ* and *mcm2-3A* mutations neutralizes single *dpb3Δ* or *mcm2-3A* mutants’ strand bias in transferring parental histone H3-H4 tetramers during DNA replication. (**A**) Procedure for monitoring the deposition of parental (H3K4Me3) and newly synthesized (H3K56Ac) histone H3 at early replication origins. (**B**) Snapshot of parental histone H3 (H3K4Me3) eSPAN read enrichment at leading and lagging strands at the early replication origin ARS1309 for wild-type (WT), *dpb3Δ, mcm2-3A*, and *dpb3Δ/mcm2-3A* strains. The sequence reads were mapped to both the Watson strand (red) and the Crick strand (green) of the reference genome. (**C-F**) *Top*: Heatmaps representing the bias ratio of parental histone H3 (H3K4Me3) eSPAN peaks for WT, *dpb3Δ, mcm2-3A*, and *dpb3Δ/mcm2-3A* strains at each of the 10 individual nucleosomes surrounding each of the 134 early DNA replication origins. Individual nucleosomes are represented by the circles at the top of the heatmaps, and their positions are indicated relative to the origin (−10 to +10). Each row represents the average log2 Watson/Crick ratio of H3K4Me3 eSPAN sequence reads at one origin. *Bottom*: Average bias ratio of parental histone H3 (H3K4Me3) eSPAN peaks for WT, *dpb3Δ, mcm2-3A*, and *dpb3Δ/mcm2-3A* strains at each of the 10 nucleosomes surrounding the 134 early replication origins.

As we have previously reported (9), parental histone H3-H4 tetramers in wild-type (WT) cells showed a slight lagging strand bias for all early S-phase replication origins (**Figs 1C-F**). The parental tetramer in the *dpb3Δ* mutant displayed lagging strand bias, whereas the *mcm2-3A* mutant displayed leading strand bias, which are consistent with previous reports (9,11). In the *dpb3Δ*/*mcm2-3A* mutant, the parental tetramer exhibited a slight lagging strand bias, similar to that of the WT cells (**Figs 1C, D**). Newly synthesized histone H3-H4 tetramers showed the reverse pattern of parental H3-H4 (**Suppl Fig 1**). These results suggest that no interaction exists between the two parental histone transfer pathways. If one pathway is defective, the other becomes predominant.

Here, we used the CRASH (Cre-reported altered states of heterochromatin) assay to monitor the transient loss of heterochromatin silencing at the *HML* locus during cell division (i.e., strains’ epigenetic instability phenotype) (25,26). We observed that the *dpb3Δ* and *mcm2-3A* double mutant shows an additive effect (**Suppl Figs 2A,B**). These results are similar to those reported previously (13). Next, we compared epigenetic instability in these mutants to that in a *Caf1* (*Δcac1*) mutant; *Caf1* is a chaperone for newly synthesized histones, and *Cac1* is its large subunit. Instability in the *Δcac1* mutant was much greater than in the parental histone chaperone mutants (**Suppl Figs 2A,B**)(26). This finding suggests that new histone assembly is more relevant to epigenetic integrity than transfer of parental histones. Taken together, our eSPAN and phenotype data indicate that the degree of parental histone strand bias is not directly correlated with epigenetic instability at *HML* locus (See more in discussion part).

**Figure 2.**
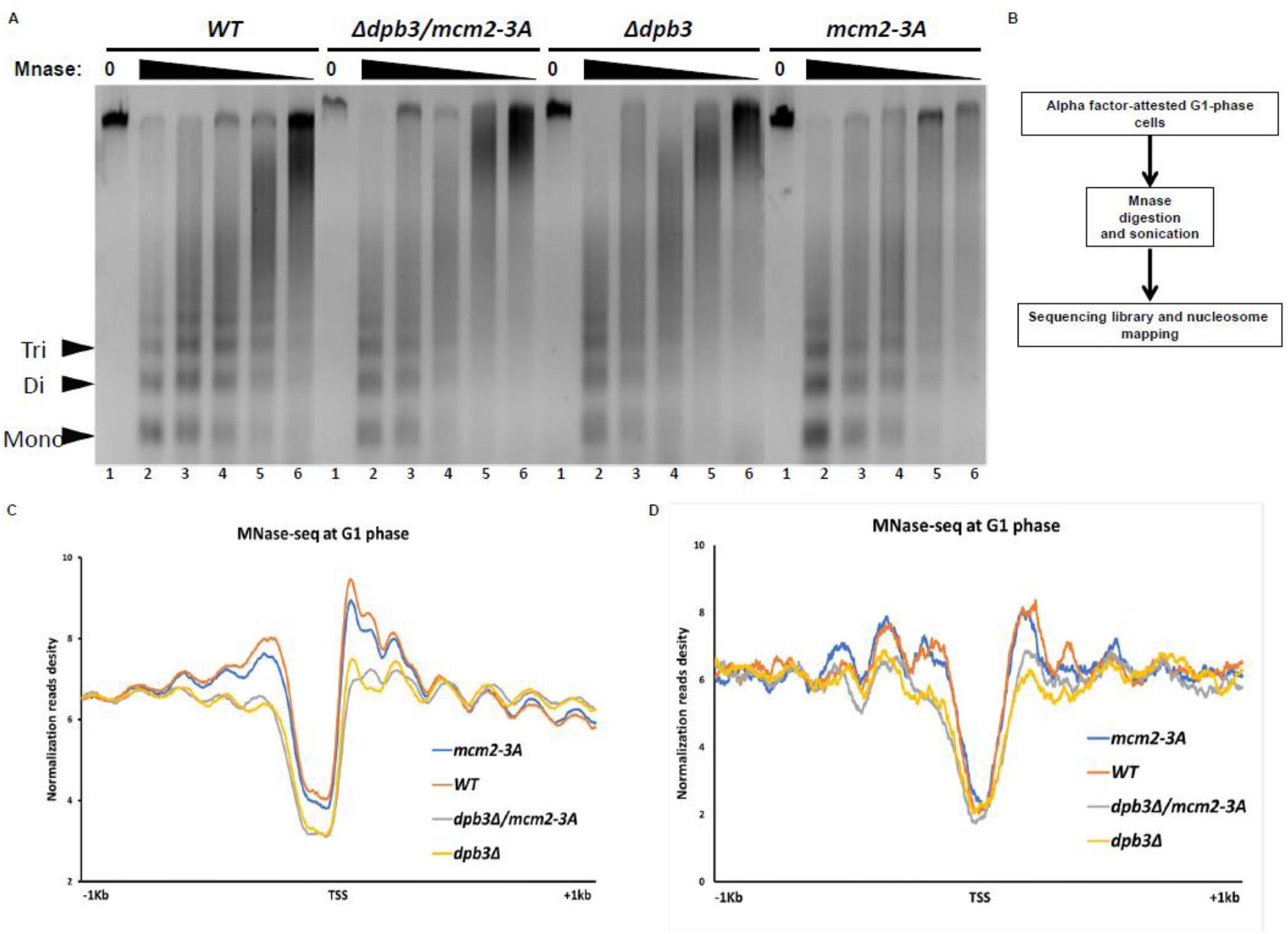
*dpb3Δ* mutations lead to subtle chromatin-structure changes. (**A**) Results from a micrococcus nuclease (MNase) chromatin accessibility assay for wild-type (WT), *dpb3Δ, mcm2-3A*, and *dpb3Δ/mcm2-3A* strains, shown on a 2% agarose gel. Chromatin sensitivity assays were performed using digestion with various concentrations of MNase for 10 min followed by quenching with stop solution and treatment with RNase. High MNase mount lead to more small fragment and clear nucleosome bands (poly, tri, di, and mononuclesomes) means strong nucleosome positioning. (**B**) Procedure used for MNase-seq, to characterize nucleosome positioning in the various strains. (**C**) MNase-seq profiles of mean nucleosome occupancy around transcription start sites (TSS) for the WT, *dpb3Δ, mcm2-3A*, and *dpb3Δ/mcm2-3A* strains. (**D**) MNase-seq profiles of mean nucleosome occupancy around early replication origin sites for the WT, *dpb3Δ, mcm2-3A*, and *dpb3Δ/mcm2-3A* strains.

### The effects of *dpb3Δ* on chromatin structure

As parental histone recycling is a fundamental process in all eukaryotic cells, we investigated whether defects in these pathways had phenotypic consequences other than the loss of silencing states. First, we examined DNA replication kinetics using fluorescence activated cell sorting (FACS). DNA replication kinetics in the *mcm2-3A* strain were equivalent to those in WT cells (**Suppl Fig 3**), as previously reported (10). The *dpb3Δ* replication was slightly slower than the WT strain. The *dpb3Δ*/*mcm2-3A* strain did not display any additive effects on DNA replication progress relative to the *dpb3Δ* strain. Of note, the delayed DNA replication progress in the *dpb3Δ* strain may not be related to its defect in a parental histone chaperone, because *Dpb3* gene also plays a regulatory role in Pol ε DNA extension and DNA repair (27).

To explore whether parental histone chaperone mutants display chromatin structural changes, we performed a micrococcal nuclease (MNase) sensitivity assay on cellular bulk chromatin. The *dpb3Δ* and *dpb3Δ*/*mcm2-3A* mutants displayed evidence of a nucleosomes positioning defect and thus greater chromatin accessibility. Relative to the WT and *mcm2-3A* strains, they did not show clear mono/di/tri nucleosome bands on an agarose gel when chromatin was treated with low concentrations of MNase (**Fig 2A**).

We carried out MNase-seq to further characterize the nucleosome positioning defect in the mutants. Cells were arrested in G1-phase with alpha factor because nucleosome positioning is strongest in this phase (**Fig 2B**). After MNase digestion and library preparation/sequencing, sequences were mapped to the all-nucleosome positions map of the whole yeast genome. The average nucleosome position profile of all genes with transcription starts sites (TSS) revealed a nucleosome-free region (NFR) and two well-positioned nucleosomes (+1 and −1 nucleosomes) surrounding the TSS (**Figure 2C**). The nucleosome positions and occupancy in the *mcm2-3A* strain were not noticeably different from those in the WT strain. However, the *dpb3Δ* and *dpb3Δ*/*mcm2-3A* strains displayed less clear positioning at the +1 and −1 nucleosomes relative to the WT strain (**Figure 2C**). MNase-seq data were also mapped to replication origins, and an NFR was also clearly observed there (**Figure 2D**). Similar to in the TSS regions, nucleosome positioning at the origins of replication were much weaker in *dpb3Δ* and *dpb3Δ*/*mcm2-3A* mutants than in the WT or *mcm2-3A* strains (**Figure 2D**).

To investigate whether the chromatin structure changes in the *dpb3Δ* mutant result in global changes to the levels of histone tail modifications or the major chromatin silencer protein Sir2, we performed western blot analysis on whole-cell protein lysates (**Suppl Fig 4A**). Levels of H3K36me3, which are associated with active gene transcription (28), were slightly lower in the *dpb3Δ, mcm2-3A* and *dpb3Δ*/*mcm2-3A* mutants than in the WT strain. This indicates that the parental histone H3-H4 tetramer transfer defect in *dpb3Δ, mcm2-3A* and *dpb3Δ*/*mcm2-3A* mutants may lead to reduced H3K36me3 in replicating cells. However, we observed no noticeable differences between the *dpb3Δ, mcm2-3A, dpb3Δ*/*mcm2-3A*, and WT strains with regard to H3K56Ac, H3K4me3, or Sir2 levels suggesting that change chromatin structure in *dpb3Δ* and *dpb3Δ*/*mcm2-3A* mutants is attributed to defect in transfer of parental histone.

### Deficiency in parental histone chaperones increases genome instability

As chromatin structure is altered in *dpb3Δ* and *dpb3Δ*/*mcm2-3A* mutants, we also tested their sensitivity to replication stress by exposing them to hydroxyurea. The *dpb3Δ*/*mcm2-3A* mutant did not show growth defects on 100 mM hydroxyurea plates (**Suppl Fig 4B**).

Next, to examine whether the parental histone chaperone mutants displayed high levels of DNA strand breaks, we used fluorescence microscopy to measure the frequency of Rad52 foci, which labels the sites of DNA lesion repair (29). The number of foci in the *dpb3Δ, mcm2-3A*, and *dpb3Δ*/*mcm2-3A* mutants were nearly twice that of WT cells (**Figs 3A,B**). These results indicate increased spontaneous DNA strand breaks in parental histone chaperone mutants.

**Figure 3.**
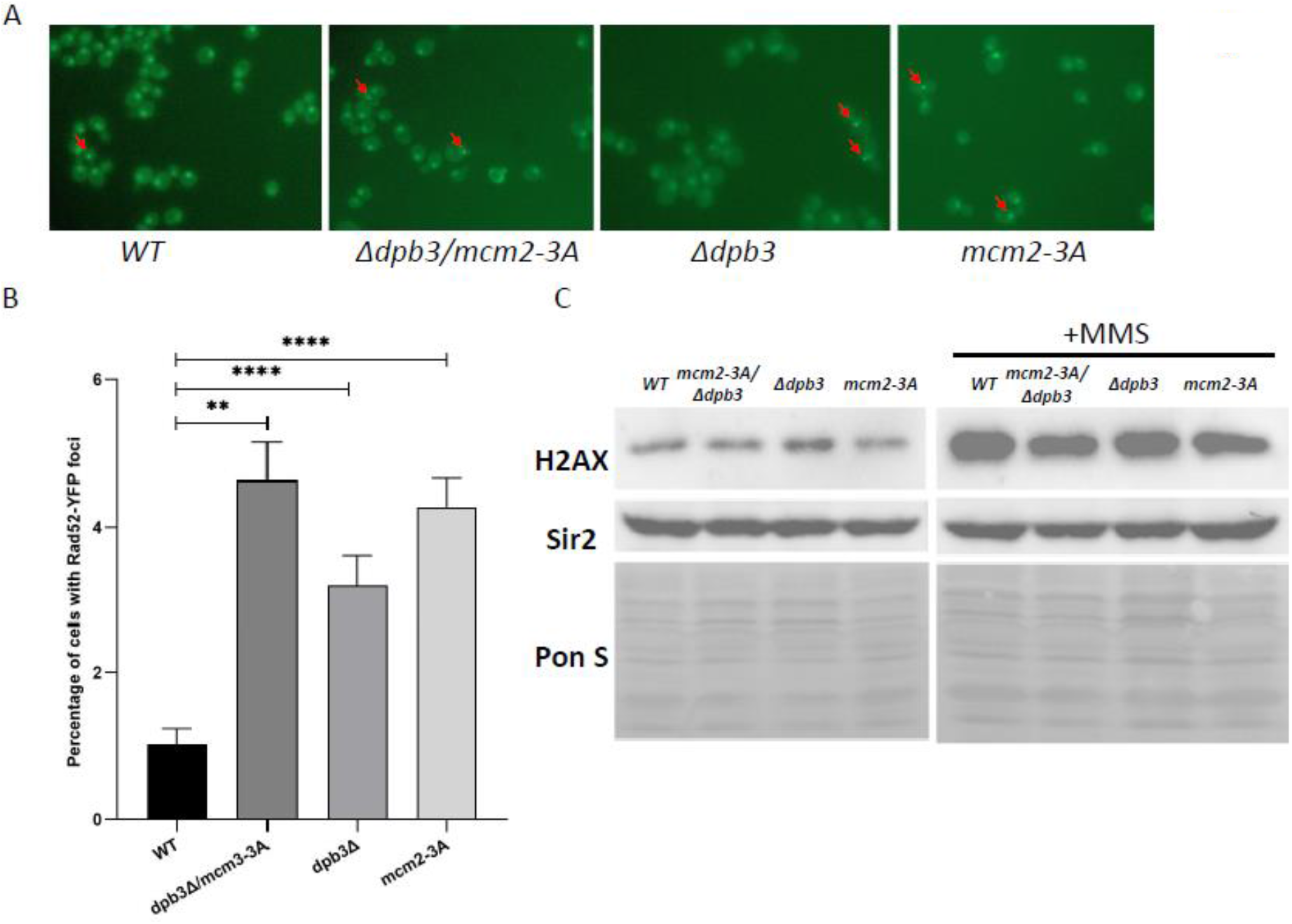
Parental histone chaperone mutants exhibit a slight increase in DNA lesions during S-phase, but a systemic DNA damage response is not triggered. (**A**) Representative fluorescence images of RAD52-YFP expression, which represent sites of DNA lesion repair, in wild-type (WT), *dpb3Δ, mcm2-3A*, and *dpb3Δ/mcm2-3A* strains. The RAD52-YFP foci are marked with red arrows. (**B**) Average percentage of cells with RAD52-YFP foci in WT, *dpb3Δ, mcm2-3A*, and *dpb3Δ/mcm2-3A* strains. Asterisks indicate statistically significance between two strains. **p < 0.01, ***p < 0.001, ****p < 0.0001. (**C**) Immunoblot analysis of H2AX levels, a marker of the DNA damage response in yeast, before and after treatment with methylmethane sulphonate (MMS; a DNA damaging agent) in WT, *dpb3Δ, mcm2-3A*, and *dpb3Δ/mcm2-3A* strains.

DNA strand breaks in yeast may activate cell cycle checkpoints and induce histone H2A SQE phosphorylation at the C terminus (SQE phosphorylation functions similar to γH2AX in mammalian cells (30). Western blot analysis showed no noticeable difference in H2A phosphorylation between the WT, *dpb3Δ, mcm2-3A*, or *dpb3Δ*/*mcm2-3A* strains (**Fig 3C**). Next, we measured H2A phosphorylation after exposing the strains to methyl methanesulfonate (MMS), which damages DNA and can induce cellular H2A phosphorylation (30), to determine whether our H2A phosphorylation antibody was effective. MMS greatly increased H2A phosphorylation levels (**Fig 3C**), as expected. These findings indicate that parental histone chaperone mutations increase DNA strand lesions but do not trigger a systemic DNA damage response.

### Spontaneous homologous recombination decreases in parental histone chaperone mutants

We hypothesized that histones in H3-H4 tetramers that dissociate from DNA in the parental histone chaperone mutants are released as soluble (free) proteins. To test this possibility, we analyzed the amount of non-chromatin-associated H3 during late S-phase. A significant increase in soluble H3 was observed in *dpb3Δ, mcm2-3A*, and *dpb3Δ*/*mcm2-3A* mutants relative to WT cells (**Fig 4B**). More free H3K4me3 was also observed in *dpb3Δ*/*mcm2-3A* mutants than in WT cells. Interestingly, total H3K4me3 levels were lower in the *dpb3Δ* and *dpb3Δ*/*mcm2-3A* mutants than in the WT or *mcm2-3A* strains in S-phase (**Fig 4B**) but not in unsynchronized log phase (**Suppl Fig 4A**). This difference may be due to the slower progress of DNA replication and parental histone recycling in *dpb3Δ* mutants.

**Figure 4.**
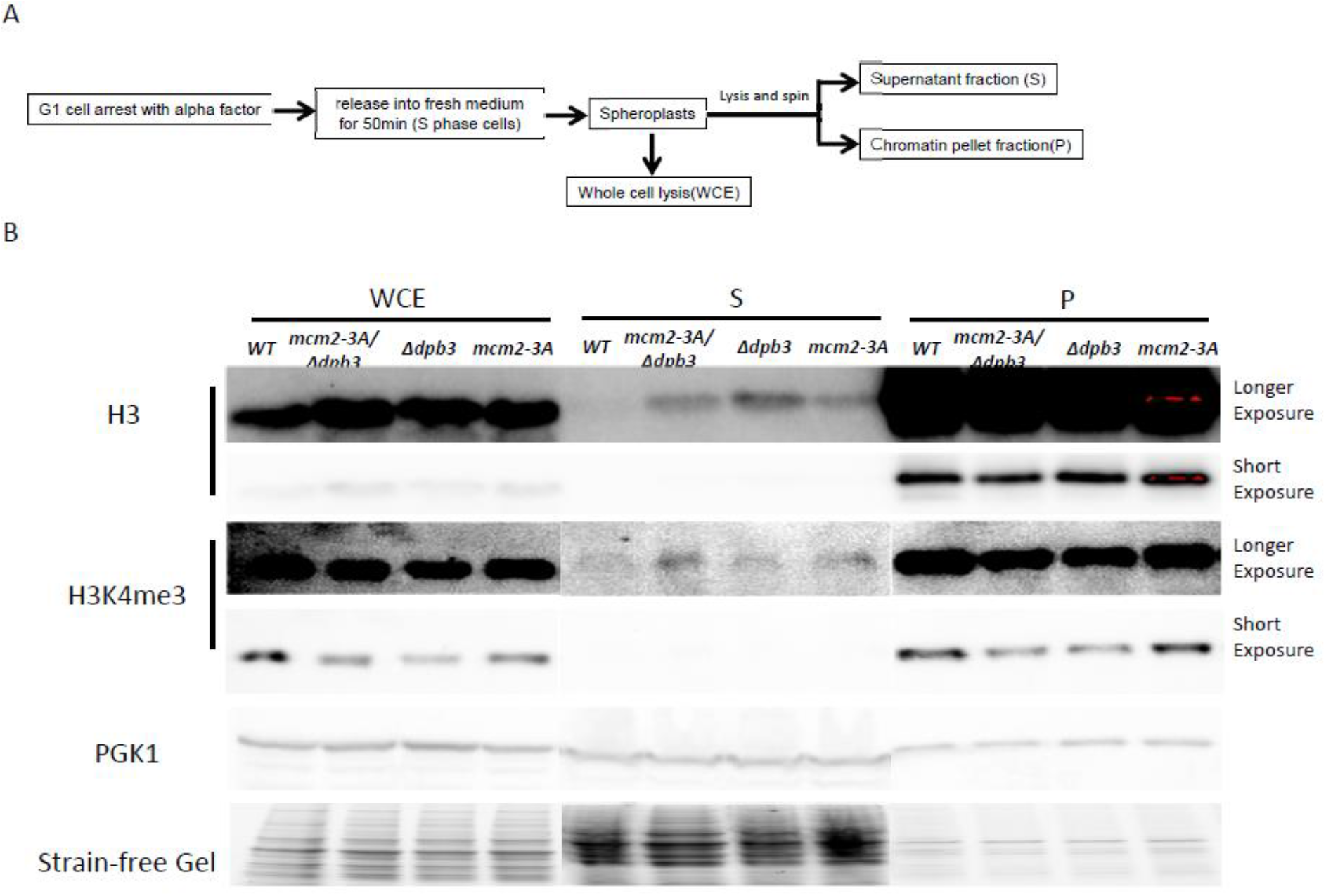
Parental histone chaperone mutations increase free histone levels in the cell. (A) Procedure to monitor free histone levels inside wild-type (WT), *dpb3Δ, mcm2-3A*, and *dpb3Δ/mcm2-3A* strains. (B) Immunoblot of H3 and H3K4me3 (a marker for parental histone H3) in the whole cell extract, soluble fraction, and chromatin fraction of WT, *dpb3Δ, mcm2-3A*, and *dpb3Δ/mcm2-3A* strains.

A previous report suggested that decreasing free histone levels by deleting one of two histone H3-H4 gene copies increases the HR rate by promoting HR factors’ binding to DNA strand breaks (31). Based on these data and the higher levels of soluble histones observed in the *dpb3Δ, mcm2-3A*, and *dpb3Δ*/*mcm2-3A* mutants, we hypothesized that the mutants might also display a decreased HR rate. To test this hypothesis, we performed a direct yeast transformation assay in which HR could be measured by neomycin and fluoroorotic acid (FOA) resistance (**Fig 5A**). The *dpb3Δ* and *mcm2-3A* mutants showed a nearly 70% reduction in HR compared to the WT strain, and the *dpb3Δ*/*mcm2-3A* mutant showed less than 10% the HR frequency of the WT strain (**Fig 5B**). We did not recover any resistant colonies in the *rad52Δ* mutant used as a negative control (data not shown).

**Figure 5.**
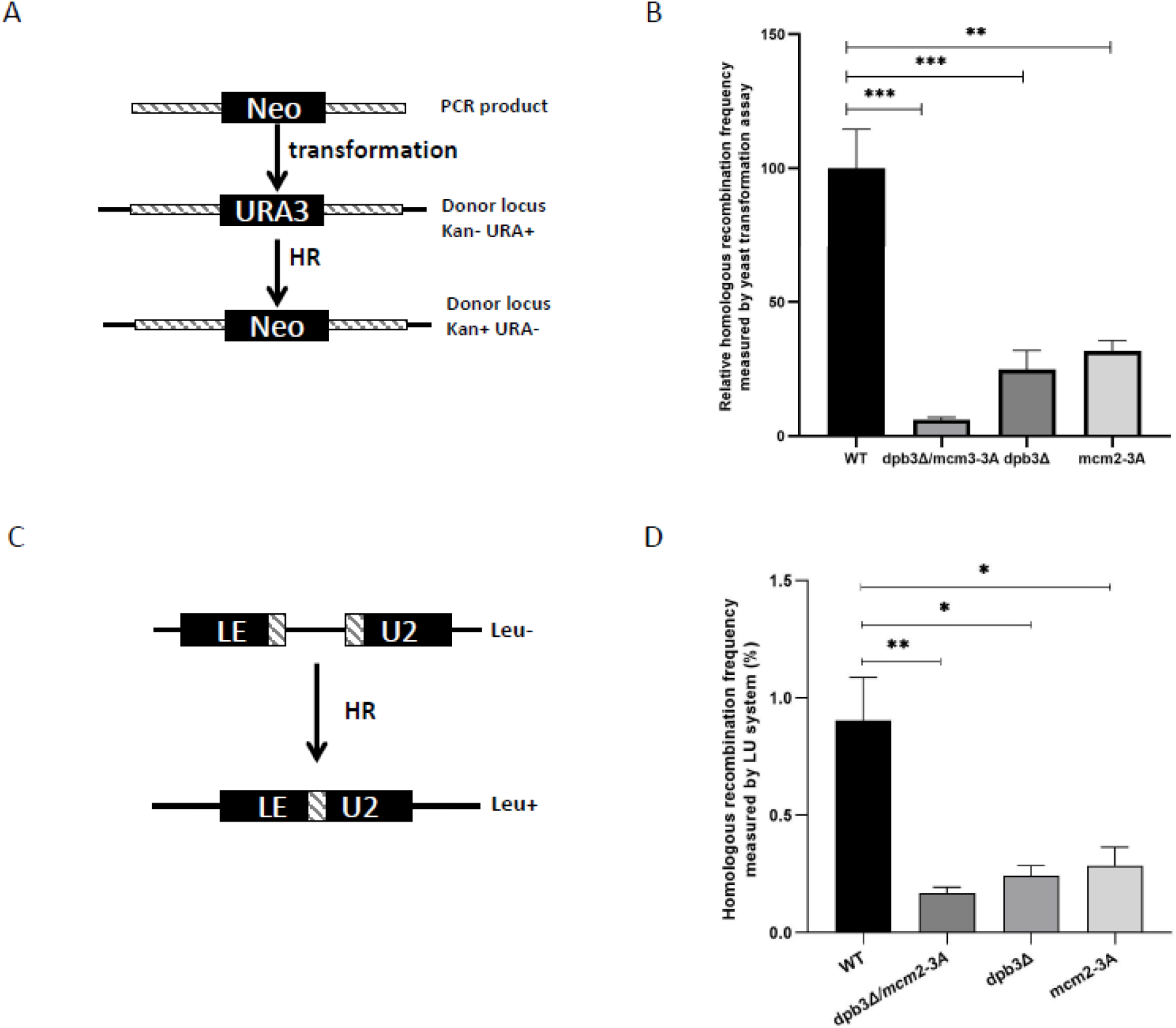
Parental histone chaperone mutations decrease the frequency of homologous recombination. (A) Procedure for quantifying HR efficiency in wild-type (WT), *dpb3Δ, mcm2-3A*, and *dpb3Δ/mcm2-3A* strains transformed with the *Neo* gene flanked by the 750-bp 5’ UTR and 550-bp 3’ UTR of the *URA3* gene. (B) Relative HR frequency of *dpb3Δ, mcm2-3A*, and *dpb3Δ/mcm2-3A* mutants relative to the WT strain. Asterisks indicate statistically significant difference between two strains. **p < 0.01, ***p < 0.001. (C) Procedure for quantifying HR efficiency using the LU system in WT, *dpb3Δ, mcm2-3A*, and *dpb3Δ/mcm2-3A* strains. (D) Percentage of WT, *dpb3Δ, mcm2-3A*, and *dpb3Δ/mcm2-3A* cells undergoing HR, as detected using the LU system. Asterisks indicate statistically significance between two strains. *p < 0.05, **p < 0.01.

To exclude the possibility that these results could reflect variation in the amount of DNA transformed into the cells, we also quantified HR using an in vivo local homologous recombination assay system (LU), in which HR is indicated by a functional Leu2 gene (**Fig 5C**) (32). Using this system, we found the frequency of HR was greatly reduced in the *dpb3Δ* (∼75%), *mcm2-3A* (70%), and *dpb3Δ*/*mcm2-3A* (∼80%) mutants. Overall, these results support our hypothesis that higher levels of free histones, caused by mutations in parental histone H3-H4 chaperones, decrease HR frequency (**Fig 6**). Of note, cells with lower rates of HR tend to employ the NHEJ pathway, which generally induces more DNA mutations than HR and is detrimental to cells (33).

**Figure 6.**
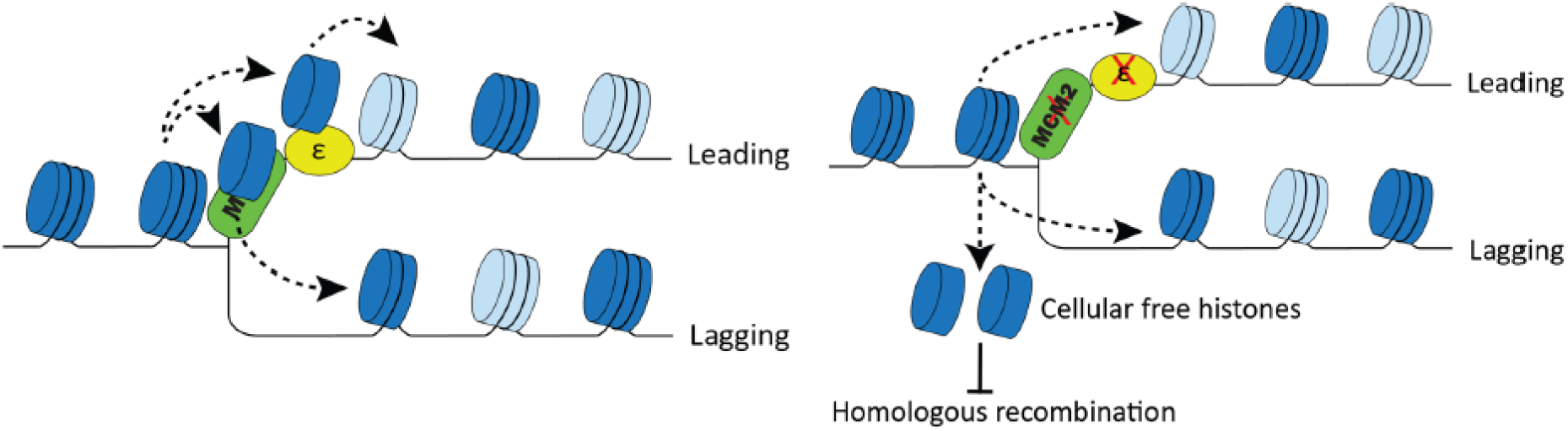
Proposed model showing that defective parental histone transfer causes increase in free histones, leading to impaired homologous recombination. In the wild type cells (left panel), parental histone H3-H4 (dark blue) can be efficiently transfer to newly synthesized DNA mediated through Polymerase ε subunit *Dpb3/Dpb4* and *Mcm2-Ctf4-Polα* axis following DNA replication fork. The new histone H3-H4 (light blue) is deposited following the empty space left by the parental histone. In the *dpb3Δ/mcm2-3a* mutant (right panel), a small fraction of parental histone H3-H4 leave chromatin and release into free H3-H4. The higher free histone level will inhibit HR through inhibiting ssDNA resections and host HR factors (31).

## Discussion

The purpose of this study was to elucidate the role of the two parental histone transfer pathways by comparing a yeast strain with mutations in both *Dpb3* and *Mcm2* to single *Dpb3* and *Mcm2* mutants, as well as the WT strain. We performed a variety of experiments to determine the effect of disrupting each pathway upon the strand bias of parental histone transfer, chromatin structure, genomic instability, and HR. Our findings consistently showed that the proper transfer of parental histones to the leading and lagging strands of DNA during replication is required to maintain chromatin structure and genome integrity.

### Parental histone transfer and epigenetic memory

During cell division, epigenetic patterns, including histone tail modifications, must be copied to maintain proper gene expression. DNA sequence–dependent mechanisms apparently play a role in the duplication of histone tail modifications. For example, a silencer DNA sequence is located at the silenced mating type locus in budding yeast. When the silencer sequence is deleted, the silenced gene is expressed after cell division (34). In addition to the sequence-dependent gene silencing mechanism, a “copy and paste” mechanism has been proposed to explain epigenetic inheritance (35). The PRC2 complex, a major gene silencer, can bind to the histone tail modification H3K27me3 and promote further H3K27me3 modification locally (36). Similarly, in the absence of demethylase, Clr4 can perform this reader-writer function to copy H3K9me3 modifications in fission yeast (35).

In budding yeast, the major mechanism responsible for gene silencing involves histone deacetylases, such as Sir2 and others sir proteins (34). In our study, the *mcm2-3a* and *dpb3Δ* parental histone chaperone mutants showed strong strand bias in parental histone transfer during DNA replication, a striking contrast from the almost total lack of bias in the WT strain. If a “copy and paste” mechanism plays a key role in silencing at the *HML* locus in budding yeast, we would expect to observe a dramatic loss of silencing in these two mutants. However, our CRASH assay results showed the loss of silencing in both mutants was only moderate and much weaker than that in the new histone chaperone *Caf1* mutant (*cac1Δ*). Like the *mcm2-3a* and *dpb3Δ* mutants, the *dpb3Δ/mcm2-3A* mutant showed little strand bias with regard to parental histone H3-H4 tetramer transfer. However, as reported previously (13), this double mutant showed a stronger loss of silencing than either of the single mutants. Collectively, these data do not support the existence of a copy and paste mechanism in budding yeast. In this study, we only investigated silencing at the *HML* locus; whether our findings hold across the genome, such as in sub-telomeric regions, or in other organisms needs further testing.

### Free histone levels in parental histone chaperone mutants

In this study, we found that the frequency of HR in parental histone chaperone mutants (*mcm2-3a, dpb3Δ*) was significantly lower than in the WT strain. The *dpb3Δ/mcm2-3A* mutant showed an additive defect. Our biochemical analysis revealed that the underlying mechanism involved high levels of free histones in the mutants, likely released from chromatin. Recently, another research group reported that the *dpb3Δ* mutation can partially rescue the *ctf4Δ* mutant’s greater sensitivity to the DNA damaging agent methylmethane sulphonate (MMS) (23). They also observed that the frequency of template-switch copying (HR dependent) was lower in *ctf4Δ* mutant. With the additional *dpb3Δ* mutation, the template-switch was still defective and the frequency of translesion synthesis (HR independent) was higher. These observations are consistent with our findings that parental histone chaperone mutants decrease HR. The same research group showed that *mcm2-3A* and *pol1-2A* deletions, both of which affect parental histone transfer to the lagging DNA strand, were not able to rescue the MMS-sensitivity phenotype of the *ctf4Δ* mutant. This finding is consistent with our previous work (11), which showed that *Mcm2, Ctf4*, and *Pol1* function in the same pathway to mediate parental histone transfer to the lagging strand. Thus, combining these mutations (*mcm2-3A, Ctf4Δ*, and *Pol1-2A*) would not be expected to further decrease HR frequency.

It is interesting to know the impact of the mutation of newly synthesized histone H3-H4 chaperones on the frequency of HR. The newly synthesized histone chaperones *Caf1* and *Asf1* promote HR through nucleosome assembly (21,22). In human cell lines, immunofluorescence staining shows that *Caf1* is recruited to DNA double strand breaks (18). Paradoxically, in plants, *Caf1* mutants show an increased frequency of HR, probably due to greater DNA accessibility (17). Our results in budding yeast show that parental histone chaperones promote HR, but the mechanism responsible appears to involve the level of free histones in the cell rather than nucleosome assembly. A recent in vitro chromatin replication study showed that higher levels of free histones decrease the efficiency of parental histone transfer (37). As we observed higher free histone levels in parental histone chaperone mutants, the efficiency of local parental histone transfer would be predicted to fall in these strains as well. The loss of silencing we observed at the *HML* locus in parental histone chaperone mutants is consistent with this prediction although it may directly be due to the defect in parental histone chaperones.

## Methods

### Yeast strains

All *S. cerevisiae* yeast strains were of the W303-1A genetic background (MATa leu2-3 112 trp1-1 ura3-1 his3-11 ade2-1 can1-100) and are listed in **Table S1**. To create the strains, deletions, tagging, and mutagenesis were performed using polymerase chain reaction (PCR)– based methods or CRISPR-Cas9, as previously reported (11,38,39). The single *mcm2-3A* mutant has three-point mutations in the histone H3 binding domain, which disrupt the ability of parental histone H3 to bind to replicating DNA strands (10).

### eSPAN

We have previously used eSPAN to detect binding patterns of a targeted protein on leading or lagging DNA strands (24,40,41). In this study, we used H3K4me3, one of the most abundant histone methylation marks, to track parental histones during S-phase, because histone methylation of newly formed nucleosomes gradually accumulates during late S/G2-phase (9,11,42). We used H3K56Ac to track newly synthesized histone H3 in S-phase, because in S-phase it is present only on newly synthesized histones; it is completely removed during G2-phase (43,44).

We performed eSPAN following a previously described procedure (40,42). WT (cyc560), *mcm2-3A* (cyc552), *dpb3Δ* (cyc604), and *dpb3Δ/mcm2-3A* (cyc602) strains were used. Specifically, yeast cells were grown in YPD medium (1% yeast extract, 2% peptone, 2% glucose) to exponential growth phase. Cells were then arrested in G1-phase using two doses of α factor (5 µg/ml; EZBiolab) for three hours at 25°C. A small fraction of G1-sample sample was collected for MNase-seq and majority of sample was released into fresh YPD medium containing 400 mg/L BrdU and 200 mM hydroxyurea for 45 min at 30°C. Hydroxyurea stalls the replication fork but does not interfere with the process of transferring newly synthesized and parental histones (9,11).

Cells were fixed by adding freshly prepared paraformaldehyde (to 1%) at 25°C for 20 min, followed by quenching with 0.125 M glycine for 5 min at room temperature. After fixation, cells were washed twice with cold water and harvested by centrifugation at 3000 rpm for 5 min. After harvesting, cells were washed and lysed in 0.1 ml ChIP lysis buffer (50 mM HEPES [pH 8.0], 150 mM NaCl, 2 mM EDTA, 1% Triton X-100, 0.1% sodium deoxycholate) with glass beads. Lysate was harvested and washed twice with NP buffer (1.6 M sorbitol, 2 mM CaCl2, 5mM MgCl, 50 mM NaCl, 14 mM β-mercaptoethanol, 10 mM Tris-HCl [pH 7.4], 0.075% NP-40, 5 mM spermidine). Chromatin was digested with MNase (LS004797,Worthington) at 37°C for 20 min into primarily di- and mononucleosomes. The digestion was terminated with 5 µl 0.5 M EDTA and 90 µl 5X ChIP lysis buffer and was kept on ice for 30 min. Cells were then lightly sonicated for three cycles (Bioraptor Pico machine, 30 sec ON/OFF) at 4°C to release chromatin fragments into solution. Soluble chromatin (eSPAN samples) was immunoprecipitated with anti-H3K4me3 antibody (ab8580 Abcam) or anti-H3K56ac antibody (44). Protein G Sepharose beads (17-0618-02, GE Healthcare) was used to recover the targeted chromatin. After washing the beads extensively, DNA from ChIP was recovered using the Chelex-100 protocol (47).

ChIP DNA was denatured by incubating it at 100°C for 5 mins and then immediately cooling it on ice for 5 min. DNA was diluted with BrdU IP buffer (1X PBS, 0.0625% Triton X-100 [v/v]). BrdU antibody (0.17 μg/ml; 555627, BD Biosciences) was added and samples were incubated at 4°C for 2 hours. Next, 20 µl Protein G beads (17-0618-02, GE Healthcare) were added to each sample and incubated for an additional hour at 4°C. The beads were extensively washed, DNA was eluted with 100 ul 1X TE buffer containing 1% SDS, and purified using a QIAGEN MinElute PCR Purification kit. ssDNA libraries were prepared using an Accel-NGS 1S Plus DNA library kit (10096, Swift Biosciences).

### Sequence mapping and data analysis

The sequence mapping, nucleosome mapping, and eSPAN analysis was performed as previously described (9,11). Briefly, the reads were mapped back to the Saccharomyces Genome Database (http://www.yeastgenome.org/) reference genome with the Bowtie2 software(45). Only paired-end reads with both ends mapped correctly were selected for continued analysis. 120-170bp DNA fragments calculated by the paired-reads were used to obtain the nucleosome occupancy by self-developed Perl programs. To calculate the eSPAN bias pattern, the forward (Watson strand) and reverse (Crick strand) reads following the reference genome were separated by self-developed Perl programs. The nucleosomes position around DNA replication origins were previously determined (46). Total eSPAN sequence reads at ±10 nucleosomes surrounding the DNA replication origins were counted for separate strands. The log2 ratio of Watson strand reads over Crick strand reads at each nucleosome position was used to obtain the average bias pattern of eSPAN.

### Analysis of silencing-loss at the *HML* locus using the CRASH assay

WT (JRY10790), *mcm2-3A* (cyc853), *dpb3Δ* (cyc756), and *dpb3Δ/mcm2-3A* (cyc777) strains were used to measure the apparent silencing-loss rate at the *HML* locus. Briefly, 10 colonies of each strain were grown separately in YPD medium to saturation, diluted to OD600 = 0.01 in YPD, and grown for 5 hours at 30°C. For the GFP positive control, cells were treated with 20 µM nicotinamide; for the RFP positive control, cells were grown in hygromycin (200 µg/ml). The apparent silencing-loss rate at the *HML* locus was calculated by dividing the number of RFP+ GFP+ cells (cells that have recently undergone Cre-mediated recombination express GFP but not RFP) by the total number of cells with the potential to lose silencing (RFP+ GFP- and RFP+ GFP+). For each colony, 50,000 events were analyzed using a BD Fortessa cytometer.

### Chromatin fractionation

We performed chromatin fractionation following a previously published procedure (**Fig 4A**) (47). WT (cyc614), *mcm2-3A* (cyc933), *dpb3Δ* (cyc931), and *dpb3Δ/mcm2-3A* (cyc929) strains were used. Cells were grown to OD600=0.5-0.6, arrested at G1-phase using two doses of α factor (5 µg/ml), grown for 3 hours at 25°C, and released in fresh YPD medium for 50 min, yielding late S-phase cells. The cells were washed with cold water and harvested by centrifuging samples at 3000 rpm for 5 min. Harvested cells were washed with spheroplast buffer (0.6M sorbitol, 24 mM Tris [pH 7.5]), suspended in the same buffer supplemented with Zymolase (0.5mg/ml, L2524, Sigma), and incubated at 30°C with gentle shaking. Spheroplasts were then washed with cold spheroplast buffer and suspended in 425 µl spheroplast buffer and 50 µl 10X lysis buffer (500 mM potassium acetate, 20 mM MgCl2, 200 mM HEPES [pH 7.5]), along with protease inhibitors (Roche). Spheroplasts were lysed by adding 20 µl 20% triton X-100 and kept on ice for 10 min. Soluble and chromatin-bound histones were separated via centrifugation at 12,000 g for 15 min at 4°C. The soluble and chromatin fractions were probed with anti-HA (12CA5, Sigma for H3 detection), anti-H3K4me3 (ab8580, Abcam), and anti-PGK1 (459250, Thermofisher) antibodies.

### Chromatin accessibility assay using MNase

MNase preferentially digests the naked DNA between nucleosomes and is commonly used to probe chromatin structure. To measure chromatin accessibility using MNase, 100 ml log phase cells were washed with cold water and harvested by centrifugation. Spheroplasts were prepared by suspending cells in spheroplast buffer (1M sorbitol, 0.5mg/ml Zymolase, 2 mM β-mercaptoethanol) at 30°C for 20 min. Spheroplasts were washed twice with sorbitol wash buffer (1M sorbitol, 1mM PMSF, 2mM β-mercaptoethanol), suspended in 600 μl spheroplast digestion buffer (1M sorbitol, 50mM NaCl, 10 mM Tris-HCl [pH 7.5], 5mM MgCl2, 1 mM CaCl2, 1mM β-mercaptoethanol, 0.075% v/v NP-40), and divided into 200 μl aliquots containing varying concentration of MNase (Worthington Biochemical) for 10 min at 37 °C. Reactions were terminated by adding 20 μl stop solution (5% SDS, 250mM EDTA) followed by proteinase K digestion for 2 hours at 55°C. DNA was extracted twice with phenol: chloroform, and RNA was degraded by treating with RNAase A for 1 hour at 37°C. DNA was suspended in TE buffer (10 mM Tris pH 8.0, 1mM EDTA), and digested nucleosomes were separated using 2% agarose gel.

### Rad52-YFP foci counting

In budding yeast, Rad52 forms spontaneous foci, predominantly during the S- and G2-phases of the cell cycle. These foci are believed to be sites of DNA lesion repair (29). WT (cyc949), *mcm2-3A* (cyc943), *dpb3Δ* (cyc945), and *dpb3Δ/mcm2-3A* (cyc947) strains were used. We counted Rad52-YFP foci in yeast strains as previously described (29,44). Briefly, 1.5 ml cultures of yeast strains expressing Rad52-YFP and grown at 25°C were harvested, washed twice with SCM-TRP, and resuspended in SCM-TRP. Living cells were then immobilized on glass slides and images were acquired using an LSM10 confocal microscope equipped with a Plan-Apochromat 40X/0.95 oil lens. All focal planes were analyzed, and cells with Rad52-YFP foci from one imaging field were counted as positive.

### Western blot to detect yeast γ-H2AX

To detect γ-H2AX, 10 ml log phase cultures of yeast grown in YPD medium with or without 0.1% methylmethane sulphonate (MMS) were harvested and washed with 20% TCA. Cell pellets were resuspended in 250 μl 20% TCA, followed by lysis with 0.5 ml glass beads. The glass beads were removed, 0.3 ml 5% TCA was added, and precipitated proteins were collected by centrifugation. Samples were transferred to 1.5 ml tubes and 0.7 ml 5% TCA was added. Cells were centrifuged at 13,000 rpm at 4°C for 10 min and supernatant was discarded. The cell pellet was washed with 100% cold ethanal and resuspended in 100 μl SDS loading buffer with 50 μl 1 M Tris pH 8.0. After 10 min at 95°C, insoluble material was removed by centrifugation and the supernatant was fractionated by SDS-PAGE. Blotting was then performed with anti-γ-H2A antibody (ab15083).

### Measuring HR frequency

To measure HR frequency, we first transformed yeast strains with a DNA fragment containing a neomycin (G418) resistance gene (*Neo*). The *Neo* flanking sequence was 100% homologous to the yeast *URA3* locus (**Fig 5A**). Successful HR generated colonies resistant to both G418 and FOA (5 –fluoroorotic acid). WT (cyc876), *mcm2-3A* (cyc892), *dpb3Δ* (cyc881), and *dpb3Δ/mcm2-3A* (SK66) strains were used. Rad52 is required for HR. Thus, we also included a *rad52Δ* mutant as a negative control.

In addition, we quantified HR using an in vivo LU system (**Fig 5C**) (32). In this system, a 2.5-kb repeat sequence is inserted into the *Leu2* gene to disrupt its function. However, HR between the repeat sequences can restore the function of Leu2. WT (YLD87), *mcm2-3A* (SK37), *dpb3Δ* (SK36), and *dpb3Δ/mcm2-3A* (SK38) strains were used. In the experiment, six independent colonies for each strain studied were picked from fresh-cultured YPD plate, resuspended in water, and plated on SC-Leu, or YPD to determine the number of Leu+, or viable colonies, respectively. The median frequency of recombination for each strain was calculated per viable cell number (determined on YPD).

## Acknowledgments

We are grateful to Dr. Akash Gunjan for kindly providing yeast strains and plasmids. We thank Jizhi Ge and Huipeng Liu for laboratory assistance. We thank Dr. Zhiguo Zhang for suggestions on this manuscripts, Dr. Junhong Han for his suggestions and help with the Rad52 foci analysis, and Dr. Kristin Harper of Harper Health & Science Communications, LLC, for editorial support. This work was supported by NIH grant R01GM130588 (to C.Y.), the Hormel Startup Fund, and the National Natural Science Foundation of China (Grant No. 32090031, 32070610) (to H.G.).

