## Supplemental files for "Defective transfer of parental histone decreases frequency of homologous recombination in budding yeast"

Running title: Efficient parental histone transfer promotes homologous recombination

**Supplemental Figure 1. Combination of *dpb3Δ* and *mcm2-3A* mutations neutralizes the strand bias of new histone H3-H4 tetramers (H3K56Ac) in single *dpb3Δ* or *mcm2-3A* mutants.**

(**A**) Snapshot of H3K56Ac ESPAN read enrichment at leading and lagging strands at early replication origin ARS1309 in wild-type (WT), *dpb3Δ*, *mcm2-3A*, and *dpb3Δ/mcm2-3A* strains. The sequence reads were mapped to both the Watson strand (red) and the Crick strand (green) of the reference genome. (**B-E**) *Top*: Heatmaps representing the bias ratio of newly synthesized histone H3 (H3K56Ac) eSPAN peaks for WT, *dpb3Δ*, *mcm2-3A*, and *dpb3Δ/mcm2-3A* strains at each of the 10 nucleosomes surrounding each of the 134 early DNA replication origins. Individual nucleosomes are represented by the circles at the top of the heatmaps, and their positions are indicated relative to the origin (−10 to +10). Each row represents the average log2 Watson/Crick ratio of H3K56Ac eSPAN sequence reads at one origin. *Bottom*: Average bias ratio of newly synthesized histone H3 (H3K56Ac) eSPAN peaks for WT, *dpb3Δ*, *mcm2-3A*, and *dpb3Δ/mcm2-3A* strains at each of the 10 nucleosomes surrounding the 134 early replication origins.

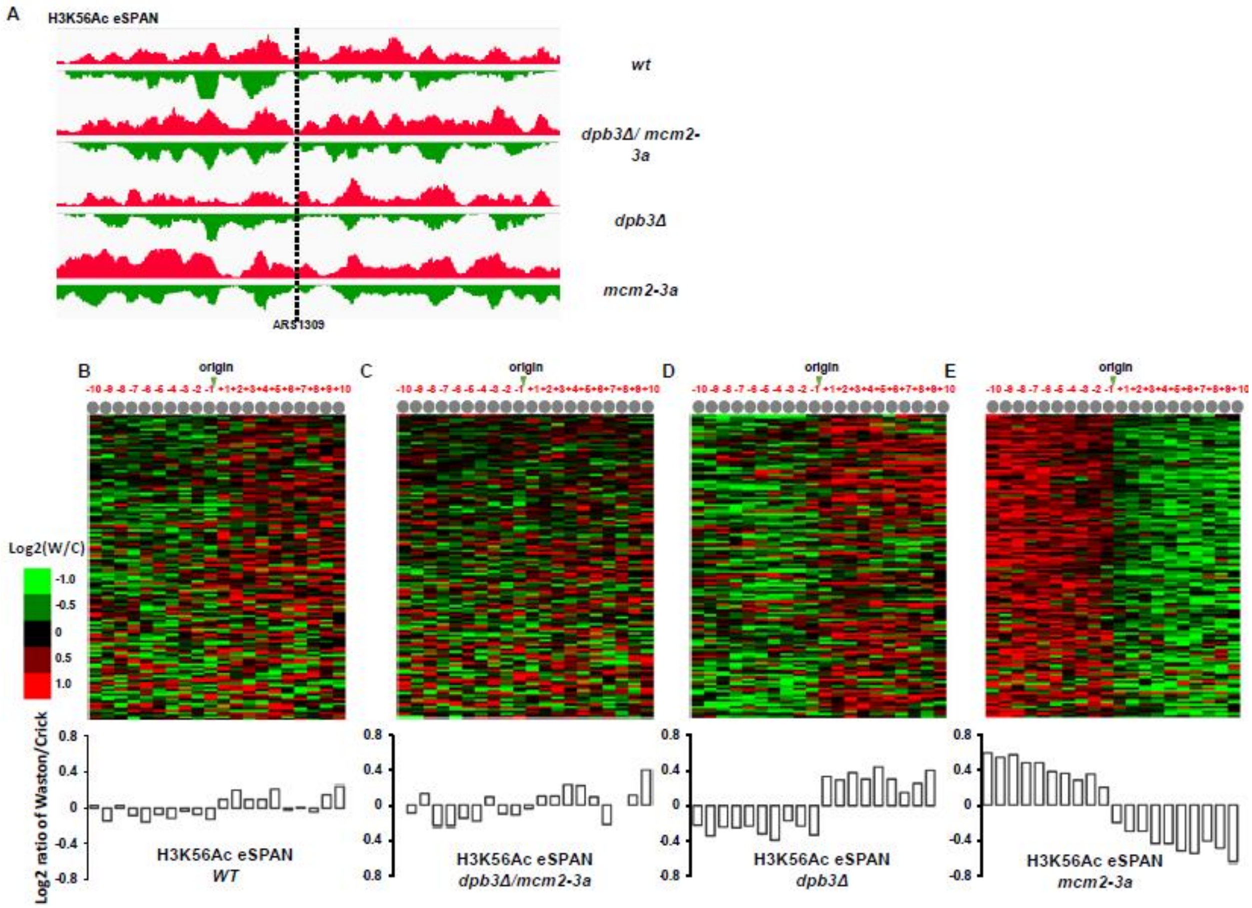

**Supplemental Figure 2. Effect of parental histone chaperone mutations on loss of silencing at the *HML* locus, as determined by CRASH assay.**

(A) Fluorescence images of colonies derived from wild-type (WT), *dpb3Δ*, *mcm2-3A*, *dpb3Δ/mcm2-3A*, and *cac1Δ* strains containing the RFP-GFP cassette at the HMLα::cre locus. The bright green sector in GFP channel or dark sector in RFP channel represent the loss of silence. (B) WT, *dpb3Δ*, *mcm2-3A*, *dpb3Δ/mcm2-3A*, and *cac1Δ* strains showed different degrees of silencing at the *HML* locus when analyzed by CRASH assay. Asterisks indicate statistically significance between two strains. *$p < 0.05$, **$p < 0.01$, ***$p<0.001$, ****$p<0.0001$.

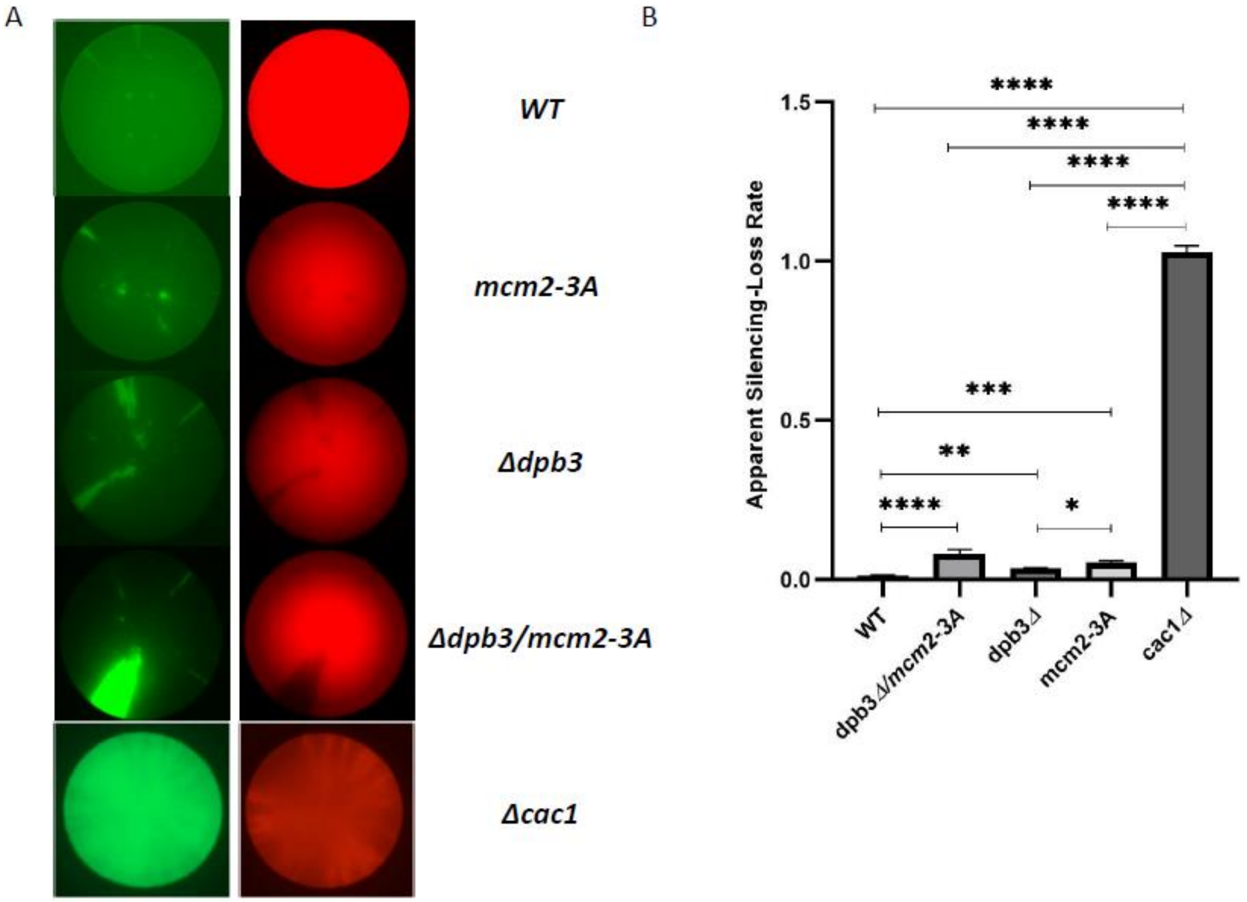

**Supplemental Figure 3. Cell-cycle progression in the parental histone chaperone mutants.**

Cell-cycle progression of the wild-type (WT), *dpb3Δ*, *mcm2-3A*, and *dpb3Δ/mcm2-3A* strains was monitored by flow cytometry after cells were released from arrest in G1-phase. A mating pheromone was added 60 min after release from G1-phase so that completion of cell division could be monitored by the disappearance of 2C cells and the reappearance of 1C cells.

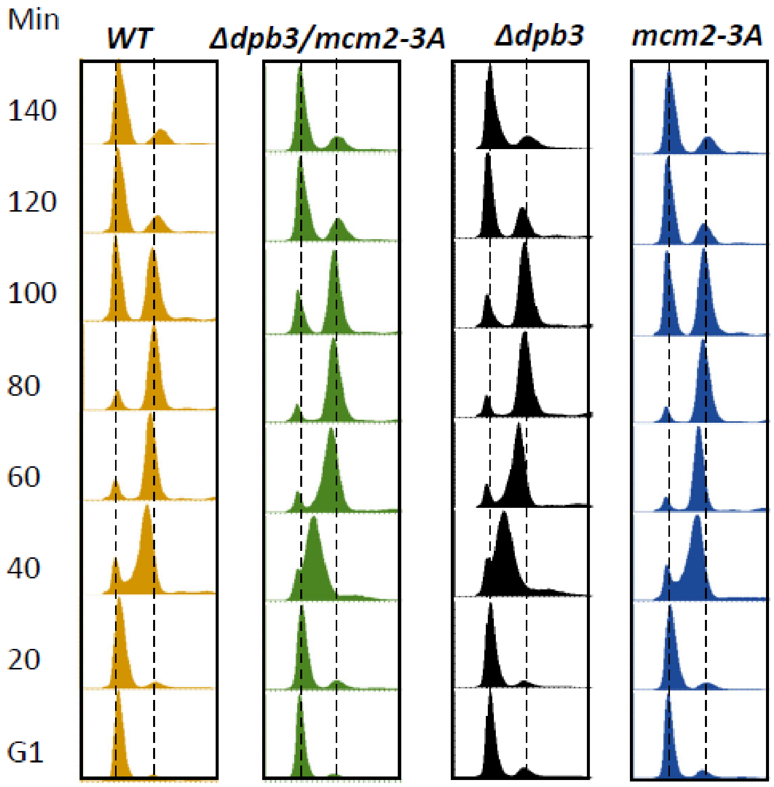

**Supplemental Figure 4. Histone modifications and cell growth in the parental histone chaperone mutants.**

(A) Immunoblot analysis of histone modification levels in whole cell extracts (WCE) from wild type (WT), *dpb3Δ*, *mcm2-3A*, and *dpb3Δ/mcm2-3A* strains. (B) Serial dilution spot assay on YPD and YPD with hydroxyurea medium for the WT and *dpb3Δ/mcm2-3A* strains.

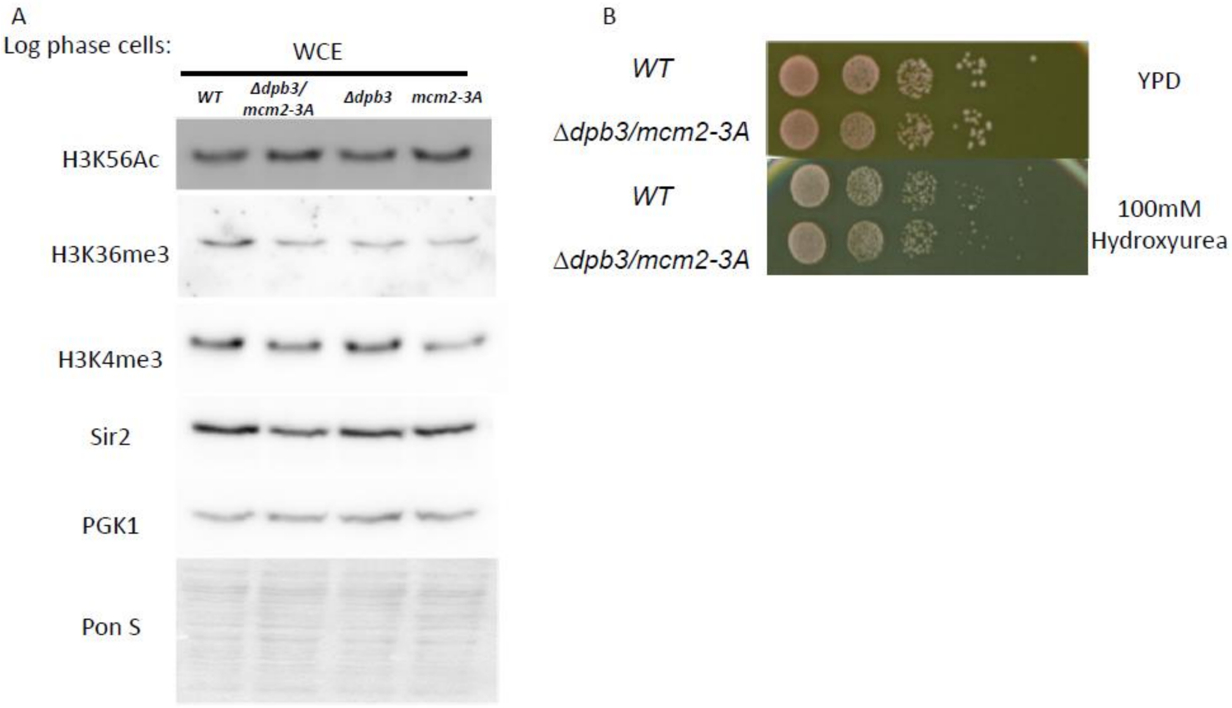

**Table S1. Yeast strains used in this study**

| **Strain** | **Genotype** | **References** |
|---|---|---|
| cyc560 | *MATA ade2-1 ura3-1 his3-11,15 trp1-1 leu2-3,112 can1-100 + URA3::BrdU–Inc* | (1,2) |
| cyc602 | *MATA ade2-1 ura3-1 his3-11,15 trp1-1 leu2-3,112 can1-100 dpb3 Δ::KanMX mcm2-3A (hphNT) + URA3::BrdU–Inc* | This study |
| cyc604 | *MATA ade2-1 ura3-1 his3-11,15 trp1-1 leu2-3,112 can1-100 dpb3 Δ::KanMX + URA3::BrdU–Inc* | (1) |
| cyc552 | *MATA ade2-1 ura3-1 his3-11,15 trp1-1 leu2-3,112 can1-100 mcm2-3A (hphNT) + URA3::BrdU–Inc* | (2) |
| JRY10790 | *MatA lys2 his3-11,15 leu2-3,112 can1-100 hml2alpha::cre ura3Δ::GPDpro-loxP-yEmRFP-CYC1term-hphMX-loxP-yEGFP-ADH1term* | (3) |
| cyc756 | *JRY10790 dpb3 Δ::NatMX* | This study |
| cyc853 | *JRY10790 mcm2-3A* | This study |
| cyc777 | *JRY10790 dpb3 Δ::NatMX mcm2-3A* | This study |
| cyc929 | *MATA ade2-1 ura3-1 his3-11,15 trp1-1 leu2-3,112 can1-100 dpb3::natR, mcm2-3A (hphNT) HHT1-3HA::kan+ URA3::BrdU–Inc* | This study |
| cyc931 | *MATA ade2-1 ura3-1 his3-11,15 trp1-1 leu2-3,112 can1-100 dpb3::natR HHT1-3HA::kan + URA3::BrdU–Inc* | This study |
| cyc933 | *MATA ade2-1 ura3-1 his3-11,15 trp1-1 leu2-3,112 can1-100 mcm2-3A (hphNT) HHT1-3HA::kan + URA3::BrdU–Inc* | This study |
| cyc614 | *MATa ade2-1 ura3-1 his3-11,15 trp1-1 leu2-3,112 can1-100 HHT1-3HA::kan + URA3::BrdU–Inc* | This study |
| cyc943 | *MATA ADE2 ura3-1 his3-11,15 trp1-1 leu2-3,112 can1-100 mcm2-3A (hphNT) Rad52-YFP+ URA3::BrdU–Inc* | This study |

| | | |
|---|---|---|
| cyc945 | *MATA ADE2 ura3-1 his3-11,15 trp1-1 leu2-3,112 can1-100 dpb3D::natR bar1 Rad52-YFP+ URA3::BrdU–Inc* | This study |
| cyc947 | *MATA ADE2 ura3-1 his3-11,15 trp1-1 leu2-3,112 can1-100 mcm2-3A (hphNT) dpb3D::natR Rad52-YFP+ URA3::BrdU–Inc* | This study |
| cyc949 | *MATA ADE2 ura3-1 his3-11,15 trp1-1 leu2-3,112 can1-100 Rad52-YFP+ URA3::BrdU–Inc* | This study |
| YLD87 | *W303 MATa his3::pRS314-LU-HIS3* | (4) |
| SK36 | *MATA dpb3::KANMX, his3::pRS314-LU-HIS3* | This study |
| SK37 | *MATA mcm2-3A HYg, his3::pRS314-LU-HIS3* | This study |
| SK38 | *MATA mcm2-3A -HYg, dpb3::KanMX his3::pRS314-LU-HIS3* | This study |
| cyc876 | *MATa ade2-1 his3-11 trp1-1 leu2-3,112 can1-100, URA3* | This study |
| cyc881 | *MATa ade2-1 his3-11 trp1-1 leu2-3,112 can1-100 dpb3::natR URA3* | This study |
| cyc892 | *MATa ade2-1 his3-11 trp1-1 leu2-3,112 can1-100 mcm2-3a URA3* | This study |
| SK66 | *MATa ade2-1 his3-11 trp1-1 leu2-3,112 can1-100 mcm2-3a dpb3::natR URA3* | This study |
